# Genetic and phylogenetic analysis of dissimilatory iodate-reducing bacteria identifies potential niches across the world’s oceans

**DOI:** 10.1101/2020.12.28.424624

**Authors:** Victor Reyes-Umana, Zachary Henning, Kristina Lee, Tyler P. Barnum, John D. Coates

## Abstract

Iodine is oxidized and reduced as part of a biogeochemical cycle that is especially pronounced in the oceans, where the element naturally concentrates. The use of oxidized iodine in the form of iodate (IO_3_^-^) as an electron acceptor by microorganisms is poorly understood. Here, we outline genetic, physiological, and ecological models for dissimilatory IO_3_^-^ reduction to iodide (I^-^) by a novel estuarine bacterium, *Denitromonas iodocrescerans* strain IR-12, *sp. nov*. Our results show that dissimilatory iodate reduction (DIR) by strain IR-12 is molybdenum-dependent and requires an IO_3_^-^ reductase (*idrA*) and likely other genes in a mobile cluster with a conserved association across known and predicted DIR microorganisms (DIRM). Based on genetic and physiological data, IO_3_^-^ is likely reduced to hypoiodous acid (HIO), which rapidly disproportionates into IO_3_^-^ and iodide (I^-^), in a respiratory pathway that provides an energy yield equivalent to that of nitrate or perchlorate respiration. Consistent with the ecological niche expected of such a metabolism, *idrA* is enriched in the metagenome sequence databases of marine sites with a specific biogeochemical signature and diminished oxygen. Taken together, these data suggest that DIRM help explain the disequilibrium of the IO_3_^-^:I^-^ concentration ratio above oxygen minimum zones and support a widespread iodine redox cycle mediated by microbiology.

## Introduction

Iodine (as ^127^I) is the heaviest stable element of biological importance and an essential component of the human diet due to its role in thyroxine biosynthesis in vertebrates^1–3^. Iodine is enriched in marine environments where it exists in several oxidation states, reaching concentrations of up to 450 nM^4^. In these environments, organisms such as kelp bioconcentrate iodine as iodide (I^-^) and produce volatile iodine species such as methyl iodide^5^. These volatile iodine species contribute to the destruction of tropospheric ozone (a major greenhouse gas) and aerosol formation at the marine boundary layer, consequently resulting in cloud formation and other local climatic effects^1,6^. Despite the global biological and geochemical importance of iodine, little is known about its biogeochemistry in the ocean^4^. For instance, the biological mechanism accounting for the unexpected chemical disequilibrium between I^-^ and iodate (IO_3_^-^) in seawater (I^-^:IO_3_^-^ disequilibrium) remains unknown^4^. At the physicochemical conditions of seawater, iodine is most stable as IO_3_^-7^, yet measurements of IO_3_^-^ and I^-^ in regions with high biological productivity (e.g., marine photic zones, kelp forests, or sediments), reveal an enrichment of the I^-^ ion beyond what can be explained through abiotic reduction^7,8^.

Among numerous explanations proposed for I^-^ enrichment, microbial IO_3_^-^ reduction is particularly compelling. The high reduction potential (IO_3_^-^/I^-^ *E_h_* = 0.72V at pH 8.1)^7,9^ makes IO_3_^-^ an ideal electron acceptor for microbial metabolism in marine environments. Early studies indicated common microorganisms such as *Escherichia coli* and *Shewanella putrefaciens*, reduce IO_3_^-^ to I^-^ ^9,10^. Subsequent studies associated this metabolism with the inadvertent activity of DMSO respiratory reductase enzymes in marine environments, along with specific enzymes (i.e., perchlorate reductase, nitrate reductase) that reduce IO_3_^-^ *in vitro* ^9,11,12^. However, there is little evidence that organisms hosting these enzymes are capable of growth by IO_3_^-^ reduction. While inadvertent IO_3_^-^ reduction might be mediated by marine bacteria possessing DMSO reductases, until recently, no definitive evidence existed that global IO_3_^-^ reduction is a microbially assisted phenomenon.

In support of a microbial role for the observed I^-^:IO_3_^-^ disequilibrium, previous studies demonstrated that at least one member each of the common marine genera *Pseudomonas* and *Shewanella* are capable of IO_3_^-^ reduction^12–14^. More recently, IO_3_^-^ reduction by *Pseudomonas* sp. strain SCT was associated with a molybdopterin oxidoreductase closely related to arsenite oxidase^14^. As part of this work, a dedicated biochemical pathway was proposed involving two peroxidases associated with a heterodimeric IO_3_^-^ reductase (Idr)^14^. The putative model proposes a four-electron transfer mediated by Idr, resulting in the production of hydrogen peroxide and hypoiodous acid^14^. Two peroxidases detoxify the hydrogen peroxide while a chlorite dismutase (Cld) homolog dismutates the hypoiodous acid into I^-^ and molecular oxygen, which is subsequently reduced by the organism^14^. The proposed pathway involving a molecular O2 intermediate is analogous to canonical microbial perchlorate respiration^15^. By contrast, Toporek *et al*.^18^ using the IO_3_^-^ respiring *Shewanella oneidensis* demonstrated the involvement of a multiheme cytochrome not found in *Pseudomonas* sp. strain SCT suggesting an alternative DIR pathway. The disparate mechanisms underscore the potential diversity of IO_3_^-^ respiratory processes. As such, identification of additional DIR microorganisms (DIRM) would clarify which genes are required for this metabolism and enable identification of IO_3_^-^ respiratory genes in metagenomes.

With this as a primary objective, we identified a novel marine DIRM, *Denitromonas iodocrescerans* strain IR-12, *sp. nov*, that obtained energy for growth by coupling IO_3_^-^ reduction to acetate oxidation. Taxonomic analysis placed this organism in the *Denitromonas* genus commonly associated with marine environments^19^. We used comparative genomics to identify the core genes involved in IO_3_^-^ respiration, which formed a distinct mobile genomic island. Reverse genetics, physiology, and comparative genomic data were used to propose a new model for DIR, with a confirmed role for a molybdopterin-dependent IO_3_^-^ reductase (IdrAB)^14^. A phylogenetic analysis was used to establish the distribution of this metabolism across the tree of life and measure the degree to which the genomic island is subject to horizontal gene transfer. Finally, metagenomic analysis identified the *idrA* gene in the Tara oceans datasets, enabling the correlation of DIR populations with ocean chemistry. These results together enabled the proposed model for the global distribution of the DIR metabolism and the ecology of the microorganisms involved.

## Results and discussion

### Isolation of *D. iodocrescerans*

*D. iodocrescerans* was isolated under anoxic conditions from estuarine sediment samples by selective enrichment followed by single colony isolation on agar plates. Analysis of the 16S rRNA indicated an axenic culture composed of a single phylotype (strain IR12) belonging to the *Denitromonas* genus in the beta proteobacteria identical to an uncultured *Denitromonas* clone from a metagenomic sample (GenBank: KF500791.1) (Figure 1A). The closest cultured relatives were *D. indolicum* strain MPKc^20^ (GenBank: AY972852.1, 99.46% similarity) and *D. aromaticus* (GenBank: AB049763.1, 99.40% similarity). Morphologically, strain IR12 is a rod-shaped motile cell 1-2 μm long and 0.5 μm diameter with a single polar flagellum (Figure 1B). Based on its phylogenetic affiliation, morphology, and metabolism (described below) we propose that strain IR12 represents a new species in the *Denitromonas* genus with the epitaph *D. iodocrescerans*.

**Figure 1:**
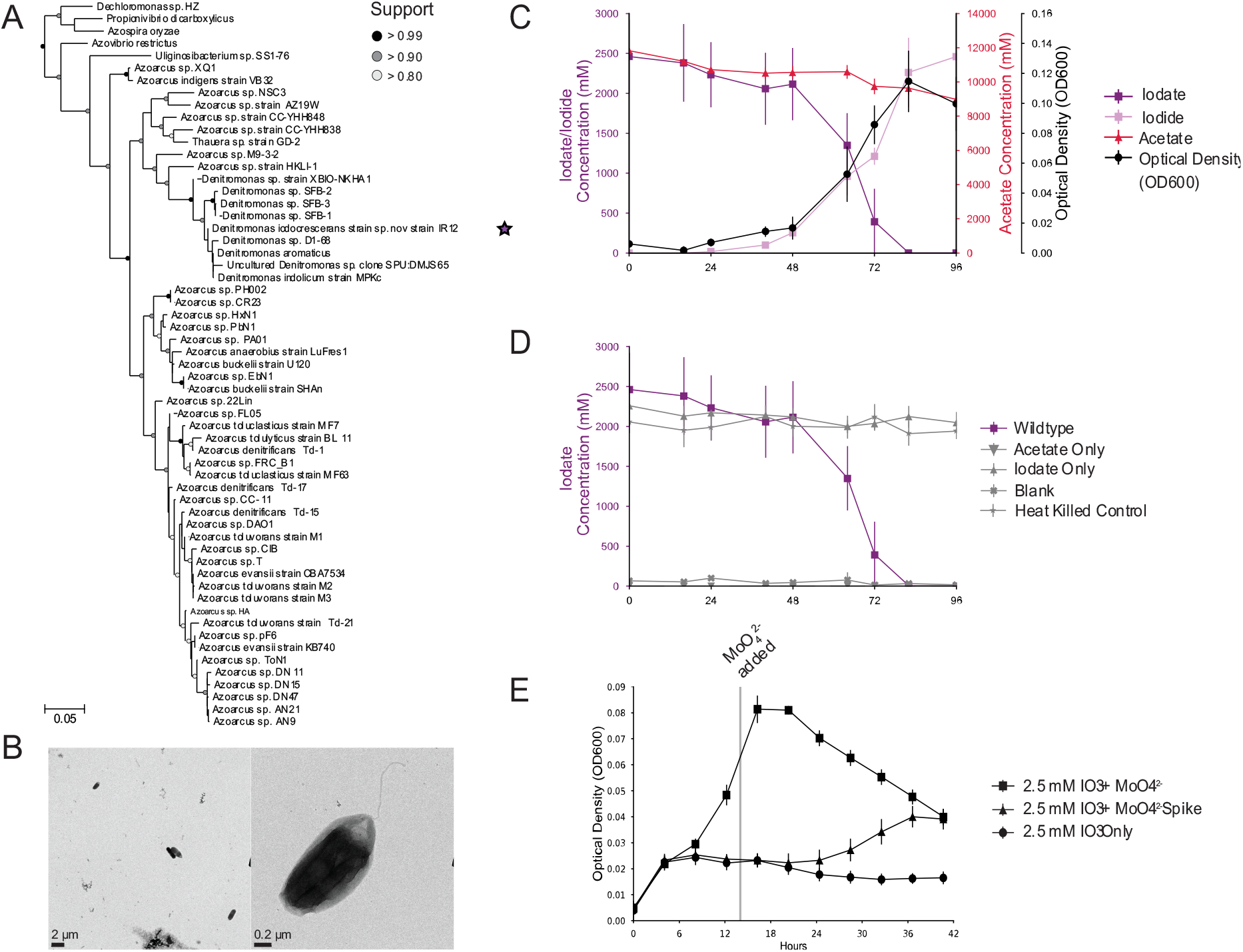
Phylogeny and Physiology of *Denitromonas iodocrescerans*. A) 16S rRNA gene phylogeny of *Denitromonas iodocrescerans* (denoted by a purple star) belonging to a subclade of *Azoarcus*, separate from other known *Azoarcus* species. B) TEM images of an active culture of *D. iodocrescerans* with the scale at 2 μm (left) and 0.2 μm (right) taken on a Technai 12 TEM. C) Iodate consumption (■), acetate consumption (▲), iodide production (■), and growth (•; measured as optical density at λ=600nm; OD600) in an active culture of *D. iodocrescerans* growing anaerobically. N=3 and error bars show standard deviation. D) Iodate consumption across all five conditions assessed in the growth experiment in C. N=3 and error bars show standard deviation. E) Optical density (OD600) in the presence (■), absence (•), and amendment after 14 hours incubation (▲) of MoO_4_^2-^. N=7 and error bars show standard deviation.

### Physiology and energetics of *D. iodocrescerans*

Cells of *D. iodocrescerans* grew on basal medium with acetate and IO_3_^-^ as the sole electron donor and acceptor, respectively (Figure 1C and D). Ion chromatography and growth studies revealed that IO_3_^-^ was quantitatively reduced to I^-^ with concomitant cell density increase. No growth or acetate consumption occurred in the absence of IO_3_^-^. Similarly, no IO_3_^-^ reduction occurred in the absence of acetate or in heat killed controls. These results indicated that IO_3_^-^ reduction was enzymatically mediated coupled to acetate oxidation and growth. Acetate-free control cultures reduced micromolar amounts of IO_3_^-^ (114 ± 34 μM, mean ± standard deviation, n=3) which was attributable to residual acetate carried over from the inoculum (**Error! Reference source not found.**). *D. iodocrescerans* consumed 2.46 ± 0.499 mM IO_3_^-^ (mean ± standard deviation, n=3) while oxidizing 2.86 ± 0.427 mM acetate (mean ± standard deviation, n=3) with a final optical density (OD600) increase of 0.109. This is equivalent to an average stoichiometry of 0.86 mol IO_3_^-^ per mol acetate. The morphological consistency between *D. iodocrescerans* and *E. coli*, suggests that an OD600 increase of 0.39 is equivalent to 1 gram of cell dry weight^21^ and that ~50% of cell dry weight is comprised of carbon^22^. Using these numbers, the corrected stoichiometry accounting for acetate incorporation into cell mass is 93% of the theoretical value according to:

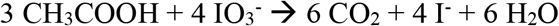

Our calculations indicate that 30.72% of total carbon is assimilated into biomass while the remaining is respired. Such a result is typical for highly oxidized electron acceptors such as oxygen, nitrate, or perchlorate^15,23^. In support of this, the calculated Gibb’s free energy and the change in enthalpy for the reduction of IO_3_^-^ per mole of electrons transferred is −115 kJ/mol e^-^ and −107 kJ/mol e^-^ respectively^24^. These values place the energy provided through IO_3_^-^ respiration akin to that of perchlorate respiration (ClO_4_^-^/Cl^-^, *E^o’^* = +0.797 V)^15^, and between that of aerobic respiration (O_2_/H_2_O, *E^o’^* = +0.820 V) and nitrate reduction (NO_3_^-^/N_2_, *E^o’^* = +0.713 V)^25^. This suggests a similar degree of carbon assimilation would be expected for IO_3_^-^ respiration ^23^.

### DIR is molybdate dependent

The reduction of oxyanions like IO_3_^-^, such as bromate, chlorate, perchlorate, and nitrate, is typically catalyzed by enzymes belonging to the DMSO reductase superfamily of molybdopterin oxidoreductases^26^. These enzymes require molybdenum as a cofactor in order to donate two electrons at a time to the receiving molecule^27^. To determine if phenotypic IO_3_^-^ reduction was molybdenum-dependent, we passaged *D. iodocrescerans* six times in aerobic, molybdate-free minimal media to remove any trace molybdenum as described in Chaudhuri *et al* ^28^. As expected, and similarly to observations with perchlorate reducing microorganisms ^28^, omitting molybdenum from the oxic medium did not affect the aerobic growth of *D. iodocrescerans* (data not shown). In contrast, no growth or IO_3_^-^ reduction was observed when these cells were passaged into molybdenum-free anoxic media with IO_3_^-^ as the electron acceptor (Figure 1E). When 0.1mM sodium molybdate was added into the non-active cultures at 14 hours post inoculation, growth and IO_3_^-^ resumed (Figure 1E). These results demonstrate that IO_3_^-^ respiration by *D. iodocrescerans* is molybdenum dependent and are consistent with the involvement of a DMSO oxidoreductase in IO_3_^-^ reduction^28^.

### Core genes required for DIR

To identify the genes required for IO_3_^-^ respiration we performed a comparative genomic analysis between the genomes of the IO_3_^-^ respiring species (*D. iodocrescerans* and *Pseudomonas* sp. SCT), and the non-IO_3_^-^ respiring close relatives (*D. halophilus* SFB-1, and *Pseudomonas* sp. CAL). Additionally, *Pseudomonas* and *Denitromonas* form phylogenetically distinct genera (*Gammaproteobacteria* and *Betaproteobacteria*, respectively), reducing the likelihood of shared gene content^29^. We surmised that DIRM must share a unique gene (or set of genes) that enables IO_3_^-^ reduction. This comparison identified 26 genes uniquely shared by the two DIRM and not found in the closely related non-IO_3_^-^ respiring species (Figure 2A; Table S2). Four of these genes were present in a gene cluster that contained genes for alpha and beta subunits of a DMSO reductase family molybdopterin enzyme related to arsenite oxidase (AioAB)^30^ supporting our result of a molybdenum dependency for this metabolism. The remaining two genes in the cluster were closely related to cytochrome C peroxidases *ccp1* and *ccp2*, possibly involved electron shuttling and oxidative stress responses^31,32^. These four genes were similar to those identified by Yamazaki *et al*. under the proposed nomenclature *idr*A, *idr*B, *idr*P_1_, *idr*P_2_ for *Pseudomonas* sp. SCT^14^ (Figure 2B). A SignalP analysis showed that *idr*P_1_ and *idr*P_2_ possessed a signal sequence for periplasmic secretion via the Sec pathway, while *idr*B used the Tat pathway^33^. By contrast *idr*A did not have a signal peptide sequence, suggesting its protein product is co-transported with IdrB into the periplasm^34^. Based on this evidence, we concluded that dissimilatory IO_3_^-^ reduction in *D. iodocrescerans* occurs entirely in the periplasm, consistent with the observation by Amachi *et al*. that associated IO_3_^-^ reductase activity in the periplasmic fractions of *Pseudomonas* strain SCT ^13^. Notably, the gene cluster lacked a quinone oxidoreductase suggesting that *D. iodocrescerans* involves the expression of a non-dedicated quinone oxidoreductase.

**Figure 2:**
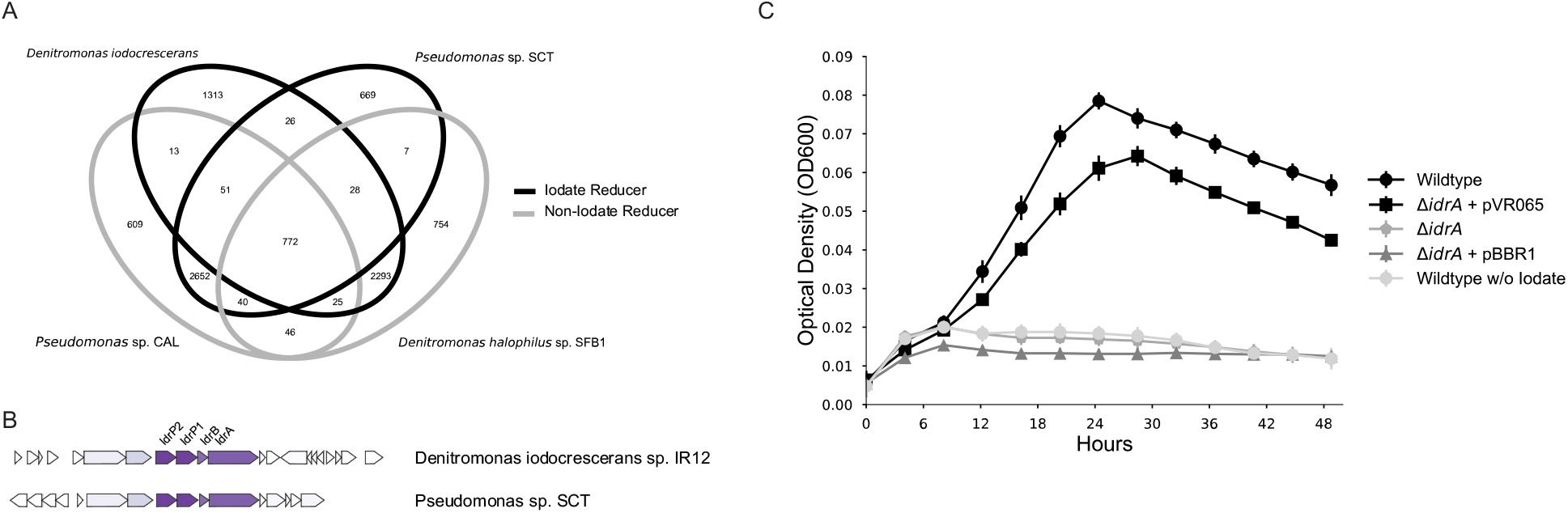
Identification of unique gene cluster in iodate reducing genomes enablling the identification and characterization of the iodate reductase (IdrA). A) A four-way comparison between two genomes from confirmed DIRM (solid line) and two genomes from closely related non-DIRM (dotted line) identifying 26 shared genes among the two taxonomically distinct iodate reducing bacteria (Table S2). B) The three genes upstream of the predicted molybdopterin oxidoreductase (IdrA) involved in DIR. C) anaerobic growth of wildtype of *D. iodocrescerans* in the presence (•) or absence 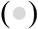 of iodate in comparison to the *DidrA mutant* 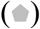 or the Δ*idrA* mutant complemented with an empty vector 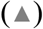 or with *idrA* in *trans* (■). N=8 and error bars represent standard deviation.

Evidence associating IdrAB to DIR, currently relies on the IO_3_^-^ consuming activity of crude cell extracts of *Pseudomonas* strain SCT and differential expression of *idrABP_1_P_2_* under IO_3_^-^ reducing conditions^14^. To validate the association between these genes and DIR in *D. iodocrescerans*, we developed a genetic system to perform targeted knockouts (see Table S1 and supplemental methods for details). The *idrA* gene was targeted since its associated molybdenum cofactor ultimately mediates the reduction of the oxyanion^26^. Upon introduction of an in-frame deletion at the *idrA* locus, the organism was incapable of growth via IO_3_^-^ respiration (Figure 2C) while growth under oxic conditions remained unimpaired. Complementation of *idrA* on a low copy number vector (pVR065) restored the IO_3_^-^ respiring phenotype demonstrating that the *idrA* gene is a prerequisite to enable IO_3_^-^ respiration (Figure 2C). Our identification of a second DIRM, in addition to *Pseudomonas* strain SCT, with an IdrAB suggests that IO_3_^-^ reduction requires a specialized molybdopterin oxidoreductase, and that other molybdopterin oxidoreductases in the genome cannot rescue the phenotype. Furthermore, our work demonstrates a distinct difference from IO_3_^-^ reduction by the multiheme cytochrome in *Shewanella* and suggests that the ability to reduce IO_3_^-^ may have evolved at least twice independently.

### An alternative DIR model

The current model for IO_3_^-^ respiration by *Pseudomonas* strain SCT proposes the donation of electrons from the quinone pool via a cytochrome c to IdrAB, to initiate reduction of IO_3_^-^ to HIO and H_2_O_2_. H_2_O_2_ is reduced to H_2_O by the peroxidases IdrP_1_ and IdrP_2_, while a chlorite dismutase (Cld)-like enzyme converts HIO to I^-^ and ½O_2_, a catalytic function that has never been demonstrated for Cld or Cld-like proteins^14^. The resultant oxygen is then further respired to H2O by a terminal oxygen reductase. The putative participation of a Cld-like protein was based on expression data rather than empirically determined activity^14^. Furthermore, comparative genomics does not support the general involvement of Cld in IO_3_^-^ respiration, as *cld* is never co-located with the IRI and is notably absent from all but two of the 145 putative DIRM genomes identified in NCBI GenBank (see below) including the genome of *D. iodocrescerans*.

Since *D. iodocrescerans* genome lacks *cld*-like genes, we propose that the primary mechanism of IO_3_^-^ respiration by this organism relies on the complex and reactive chemistry of iodine oxyanions^35^ and that the peroxidases IdrP1 and IdrP_2_ serve a critical detoxification role for inadvertent oxidants generated rather than being central components of the pathway itself. In the *D. iodocrescerans* model (F**igure 3**A), IdrAB accepts electrons from cytochrome c551, and performs a four-electron transfer, similarly to the mechanism of perchlorate reductase (Pcr)^36^, with a resultant production of the chemically unstable intermediate hypoiodous acid (HIO). This intermediate then undergoes abiotic disproportionation to yield I^-^ and IO_3_^-^ as reported in alkaline aquatic environments^16,37^, and is simplistically represented by the following equation:

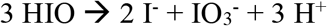

**Figure 3.**
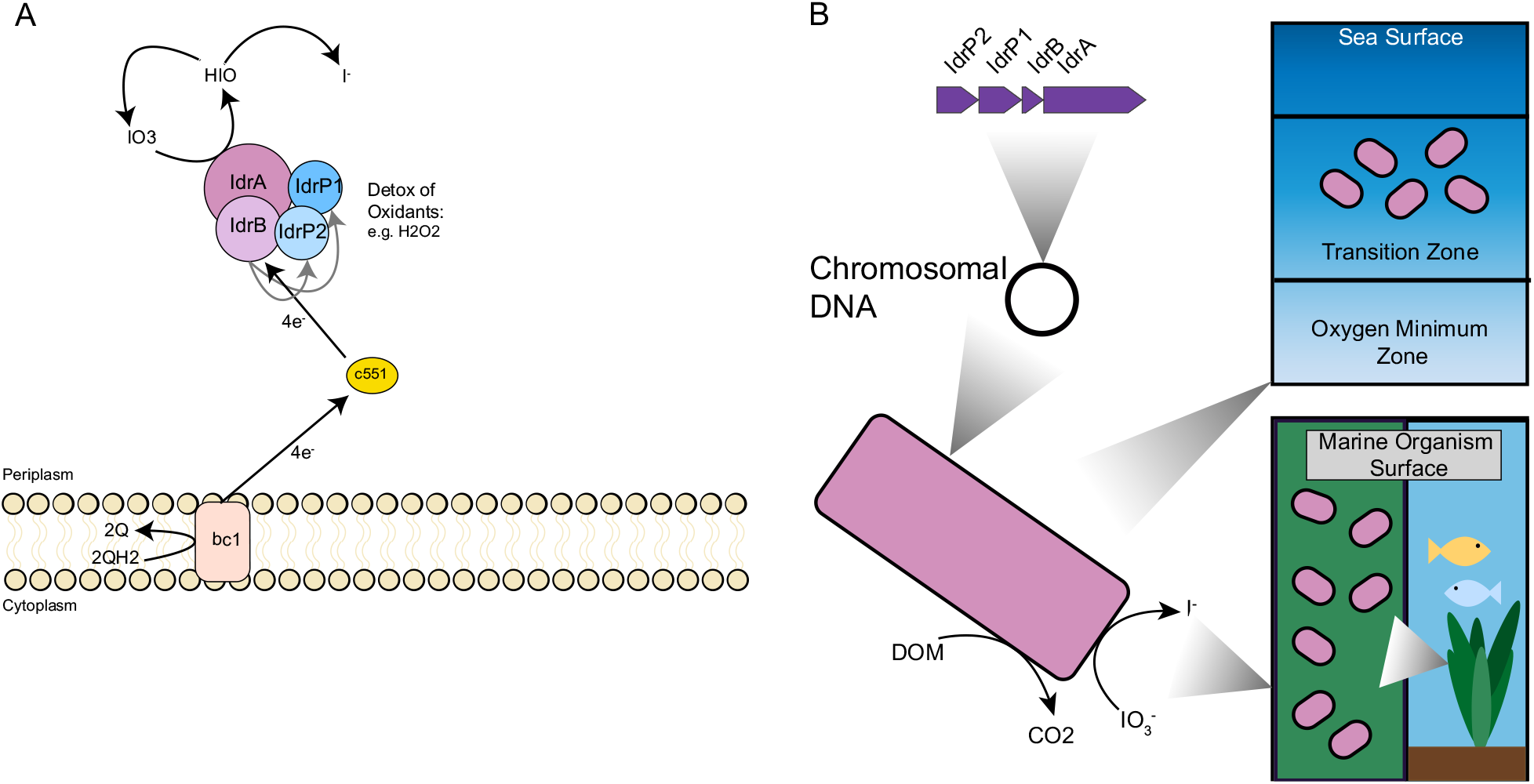
Mechanistic and Ecological Models of Iodate Reduction. A) A representation of the electron flow (black arrows) from the quinone pool to iodate in *D. iodocrescerans*. Abbreviations: QH2—reduced quinone, Q—oxidized quinone, bc1—bc1 complex, IO3^−^iodate, HIO—hypoiodous acid, I^−^iodide. Gray arrows represent micromolar production of yet unknown oxidant that is detoxified by IdrP1 and IdrP_2_. B) Ecological model of iodate reducing bacteria. Top right panel represents locations in the open ocean near oxygen minimum zones inhabited by DIORM. Bottom right panel represents host associated DIORM. DOM—dissolved organic matter.

The resultant IO_3_^-^ subsequently cycles back into the reductive pathway. In this manner, the cell completes the 6-electron reduction of IO_3_^-^ to I^-^ without invoking a Cld-like enzyme with putative capacity to dismutate IO^-^ to I^-^ and O2. This model is similar to the cryptic model for some species of perchlorate reducing microorganism which rely on the chemical reactivity of the unstable pathway intermediate chlorite (ClO2^-^) with reduced species of iron or sulfur to prevent toxic inhibition ^36,38^. We propose that the initial reduction of IO_3_^-^ at the IdrA inadvertently produces low levels of incidental toxic H2O2. This is analogous to the production of hypochlorite (ClO^-^) by respiratory perchlorate reducing microorganisms during respiration of perchlorate or chlorate^39,40^. To protect themselves from this reactive chlorine species, perchlorate respiring organisms have evolved a detoxifying mechanism based on redox cycling of a sacrificial methionine rich peptide^40^. In the *D. iodocrescerans* model for IO_3_^-^ respiration the cytochrome c peroxidases play the critical detoxification role against inadvertent H2O2 production, rather than a central role for the reductive pathway as proposed for *Pseudomonas* strain SCT^14^ (F**igure 3**A). Such a model is not only parsimonious with the predicted biochemistries and abiotic reactivities of the proteins and iodine oxyanions involved but is also consistent with the micromolar quantities of H2O2 observed by Yamazaki *et al*. during the reduction of millimolar quantities of IO_3_^-^ by *Pseudomonas* strain SCT^14^.

### Evolutionary history of DIR

Core genes for DIR were used to define the phylogenetic distribution of this metabolism. Close homologs to the catalytic subunit of IdrA were identified among genomes in NCBI GenBank. A phylogenetic tree of the DMSO reductase family (Figure 4A and 4B) confirms previous results indicating that arsenite oxidase alpha subunit (AioA) is the most closely related characterized enzyme to IdrA^14^. The extent of the IdrA clade was difficult to define because IdrA from *D. iodocrescerans* and *Pseudomonas* sp. SCT are closely related. To determine whether more IdrA homologs in this clade function as IO_3_^-^ reductases or arsenite oxidases, we performed a gene neighborhood analysis looking at the 10 genes both upstream and downstream of either the *idrA* or *aioA* locus and clustered them using MMseqs2^41^ (Figure 5). We observed a clear distinction in neighborhood synteny between genes mostly closely to *idrA* versus those most closely related to *aioA*. All neighborhoods in the *idrA* clade showed conserved synteny at *idrABP_1_P_2_* (Figure 5), whereas organisms with an AioA, showed an alternative gene structure, notably missing the cytochrome c peroxidases. Based on this pattern, all organisms possessing *idrABP_1_P_2_* genes are likely DIRM. The outgroups of IO_3_^-^ reductase in this phylogeny are homologs found in *Halorubrum* spp., which are known to oxidize arsenite^42^, and a *Dehalococcodia* bacterium (GCA_002730485.1), which also lacks the cytochrome c peroxidases in its gene neighborhood (Figure 5). Further research into these proteins may provide more information on the transition from arsenite oxidase to IO_3_^-^ reductase.

**Figure 4.**
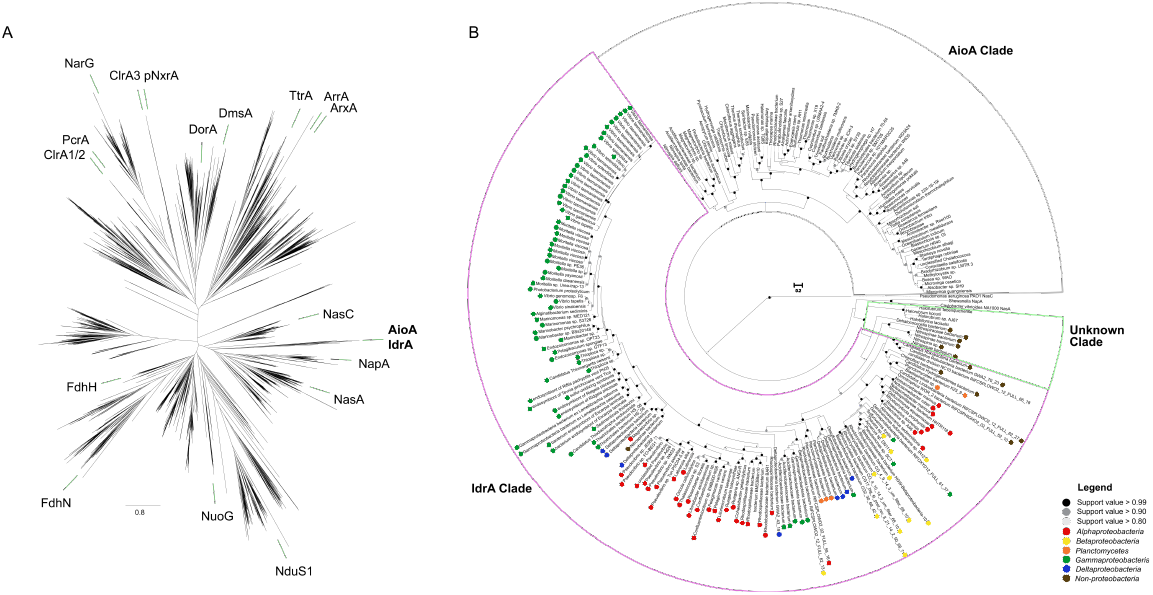
Phylogeny and Taxonomic Distribution of IdrA. A) Phylogeny of Molybdopterin Oxidoreductases (Pfam 00384) using pre-aligned proteins from the representative proteomes 55 dataset. Green bars indicate location of an individual protein in each branch belonging to the labelled group. B) Phylogeny of IdrA (purple), AioA (gray), and an unknown clade (light green) that contains proteins from organisms showing demonstrated arsenite oxidation abilities. Colored circles along the edges of the IdrA clade indicate the different Phyla each organism belongs to.

**Figure 5.**
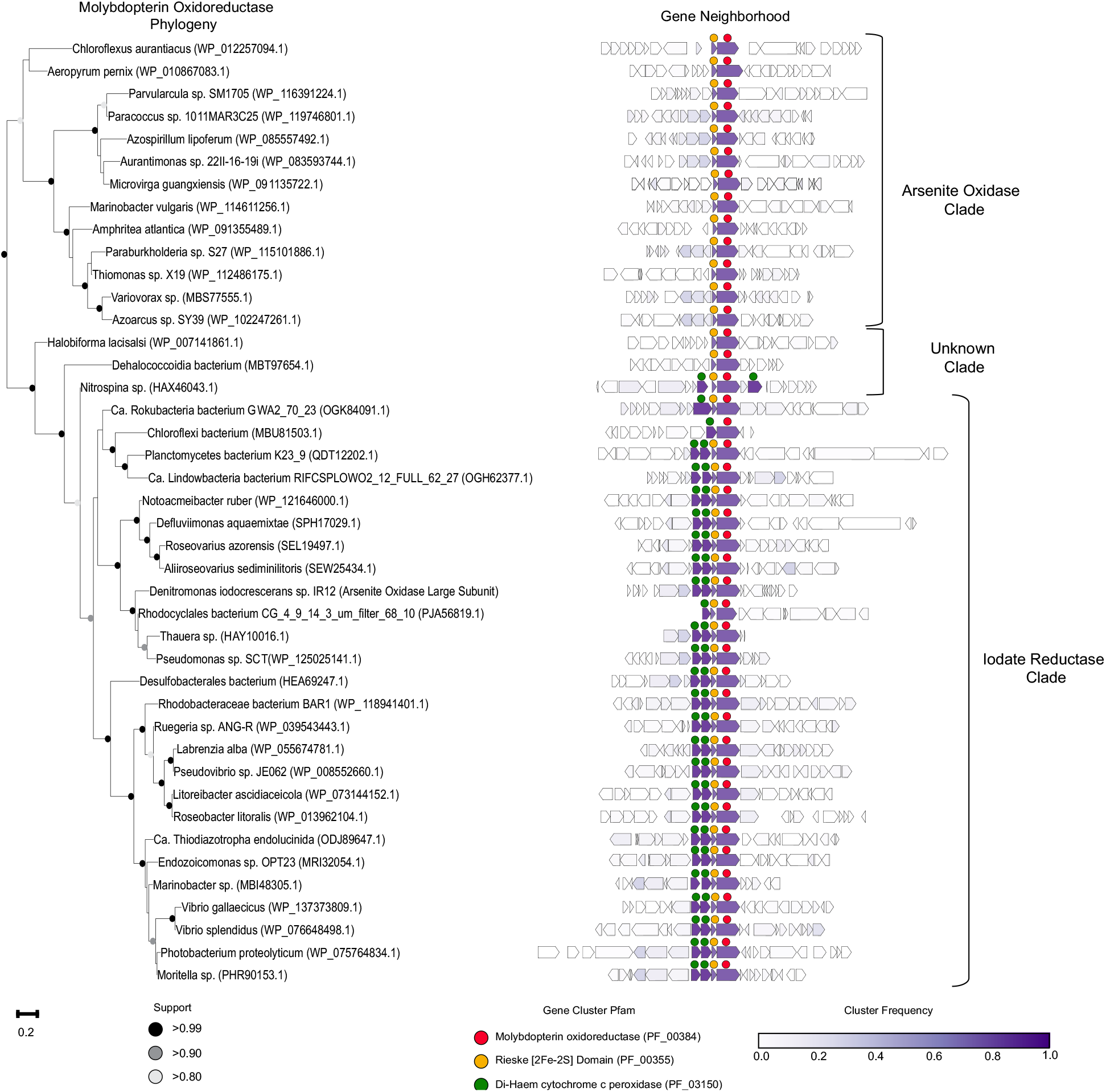
Phylogeny and Gene Neighborhoods of arsenite oxidase, iodate reductase, and the associated unknown clade. A pruned tree of the molybdopterin oxidoreductase phylogeny (left) showing a representative subset of genomes identified from Figure 3B. *Denitromonas iodocrescerans* is illustrated in bold. Genome neighborhoods (right) show 10 genes upstream and downstream (if present) from the *idrA* locus. Individual genes were clustered into groups based on amino acid similarity using MMSeqs2 and the frequency of genomes possessing an individual cluster is colored by the intensity of purple. Circles above each gene represents either the molybdopterin oxidoreductase 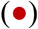, the associated Rieske containing subunit 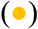, or the di-haem cytochrome c peroxidases 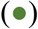.

Genes mediating IO_3_^-^ reduction were identified in 145 genomes from bacteria in the *Alphaproteobacteria*, *Betaproteobacteria*, and *Gammaproteobacteria*. Deeper branching members included members of *Planctomycetaceae* and several others belonging to the Candidate Phyla Radiation group such as, *Ca. Rokubacteria, Ca. Lindowbacteria*, and NC10 (Figure 4B)^43–45^. DIR seemed most prevalent in the phylum *Proteobacteria*, which is a pattern that has been observed for some other rare metabolisms^46^. The discordance between the taxonomy of the host organisms and the phylogeny of IdrA (Figure 4B; Figure S1)^47^ suggested that DIR is a horizontally transferred metabolism. For example, IdrA in the *Gammaproteobacterium Pseudomonas* sp. SCT was most closely related to IdrA in *Betaproteobacteria* such as *Azoarcus sp*. DN11. Additional evidence for horizontal gene transfer in individual genomes included insertion sites at the 3’ end of tRNAs, a skew in GC content, and association with other horizontally transferred genes^48,49^. In *D. iodocrescerans*, there was no significant GC skew, but we observed a tRNA^Gly^ roughly 72 kbp downstream of the *idrABP_1_P_2_* locus. While we did not detect inverted repeats, Larbig *et al*. previously demonstrated an integration site in *P. stutzeri* at tRNA^Gly50^. Additionally, numerous heavy metal resistance markers, like *mer* and *cus* genes, were found near the *idrABP_1_P_2_* locus (1.2 kbp and 22 kbp away respectively), further suggesting horizontal transfer^48,51,52^. A method to detect genomic islands in complete genomes predicted the *idrABP_1_P_2_* locus to be its own 5.8 kbp genomic island in *Azoarcus* sp. DN11, which has a complete genome and a closely related IdrA. Therefore, while there is poor conservation of genes surrounding *idrABP_1_P_2_* and questions remain about its recent evolution, the high degree of conservation of *idrABP_1_P_2_* locus itself and the phylogenetic pattern of inheritance support its description as an iodate reduction genomic island (IRI) that is subject to horizontal gene transfer. In addition to the perchlorate reduction genomic island (PRI)^46^ the IRI represents one of the few respiratory genomic islands known that crosses large phylogenetic boundaries (class, order, and family).

### Distribution of DIR populations in global oceans

Many of the organisms with genes for DIR were identified in diverse marine habitats where IO_3_^-^ reduction is suspected to occur (Table 3). For example, *Litorimicrobium taeanense* is an aerobic, non-motile, *Alphaproteobacterium* isolated from a sandy beach in Taean, South Korea^53^. Other organisms such as *Endozoicomonas sp*. OPT23 and *Litoreibacter ascidiaceicola* were isolated from marine animals such as the intertidal marine sponge (*Ophlitaspongia papilla*) and the sea squirt (*Halocynthia aurantium*), respectively^54,55^. Additionally, organisms known to accumulate iodine, such as algae^56^ are associated with these bacteria as is the case with the bacterium *Rhodophyticola porphyridii* and the red algae *Porphyridium marinum*^57^. To investigate this marine prevalence further we used the *idrA* subunit as a marker gene to determine DIRM distribution across the Tara Oceans metagenome dataset. Our approach also identified the read abundance mapping to these unique IdrA hits at the different sites by using the transcripts per million (TPM) method for read quantification^58,59^. With this method, the number of unique IdrA hits was directly proportional to the number of reads mapped to the hits (Figure 6A and 6B). In general, locations with few unique IdrA hits lacked reads mapping to IdrA (Figure 6B). We observed that 77% (74/96) of the hits arose from the mesopelagic zone at an average depth of about 461 meters (range 270m-800m) across identified stations (Figure S2). The remaining hits arose predominantly in epipelagic zones, such as the deep chlorophyll maximum in 21% of cases (20/96) and far fewer hits were observed in the mixed layer (1/96) or the surface water layer (1/96).

**Figure 6.**
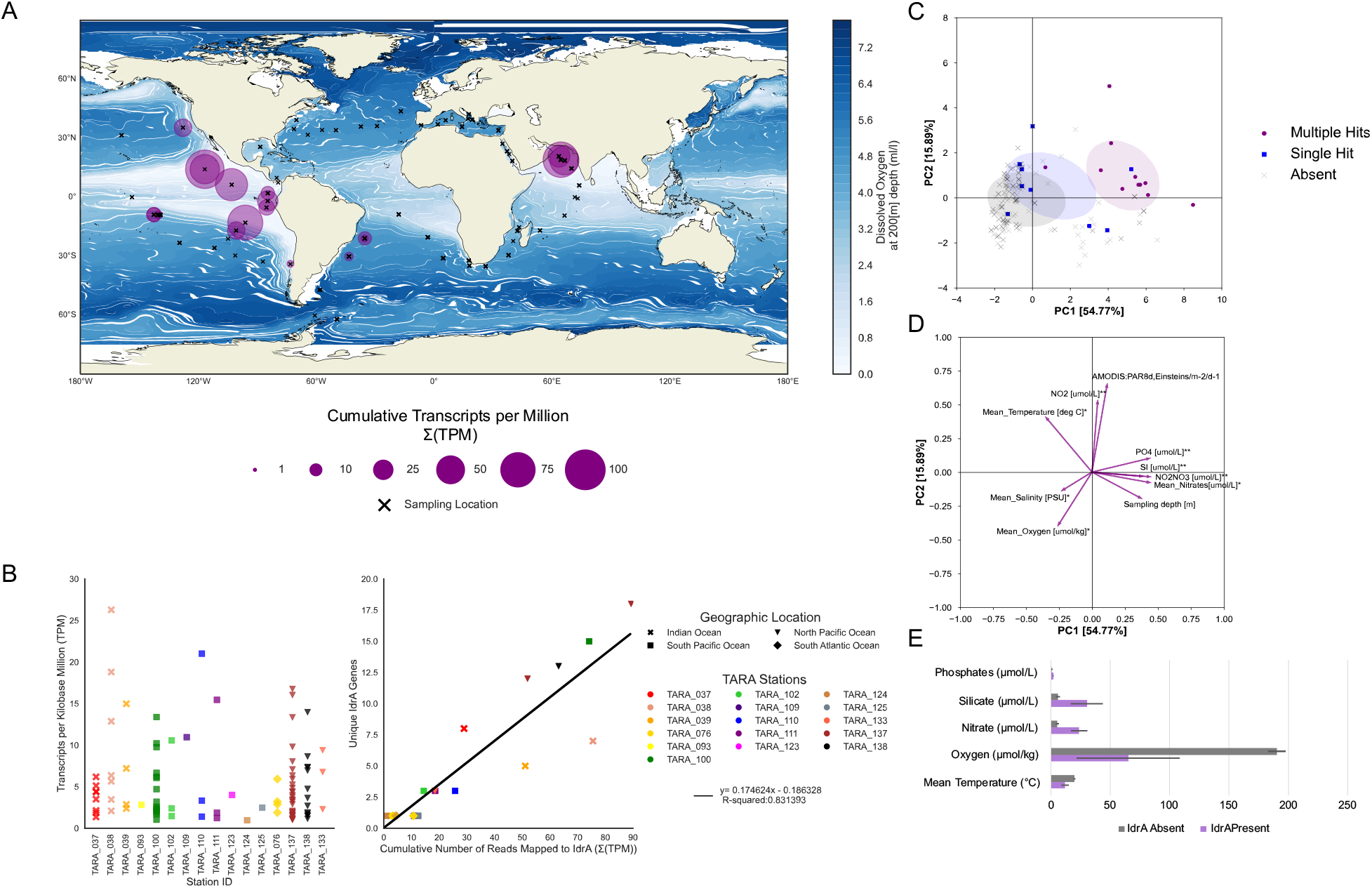
Analysis of Tara Oceans dataset identifies possible ecological niche above oxygen minimum zones. A) A map indicating sampled locations during the Tara expedition (x) alongside sampling locations with IdrA present (purple circles). Markers overlaid directly on top of each other demonstrate transect samples from different depths at a given location. Size of purple circle shows the cumulative TPM at a particular site. B) Chart on the left shows the TPM of individual hits on a scaffold organized by Tara location identifier. Coloration represents the individual Tara station while marker shape indicates general geographic location. Chart on the right correlates the number of unique IdrA hits at any given site to the cumulative TPM at an individual location. Tara station is denoted by color and general geographic location is denoted by marker shape. C) A principal component analysis displaying the first two principal components. Locations are grouped by IdrA absent 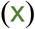, presence of a single IdrA hit 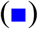, or presence of multiple hits 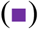. Ellipses represent 1 standard deviation of the mean. The color of the ellipse corresponds to the variable grouping. D) A loading plot of the ten variables used in the first two principal components with variables identified at the end of each arrow. E) The means of select environmental variables at IdrA present sites (purple) and IdrA absent sites (gray). Error bars indicate 95% confidence interval. Units for each of the variables are located near the variable name.

Although the presence of *idrA* exhibited some variability in depth, a geochemical feature common to all these hits was low oxygen concentrations. The vast majority of hits mapped to well-documented oxygen minimal zones in the Arabian Sea^60,61^ and the Eastern Tropical Pacific^62–64^. Similarly, the North Pacific Subtropical and Polar Front (MRGID:21484) and the North Pacific Equatorial Countercurrent provinces (MRGID:21488) are two Longhurst provinces with OMZs that stand out in the Western hemisphere. At each of these locations, the median dissolved oxygen concentration at *idrA* positive locations was consistently lower than the dissolved oxygen concentrations at *idrA* absent locations (65.24 μmol/kg versus 190.41 μmol/kg; Figure 6E). Among locations containing more than one *idrA* hit, the average oxygen concentration was about six times lower (11.03 μmol/kg); however, this average was skewed upward due to one outlier condition with 18 *idrA* hits (Cumulative TPM of 89.30; Figure S2) occurring at a dissolved oxygen concentration of 95.4 μmol/kg (TARA_137_DCM_0.22-3). Environments meeting these conditions were the most common in mesopelagic zones broadly. One notable exception were the multiple hits at the deep chlorophyll maximum (DCM) at station 137. However, further inspection of the physical environment at the DCM revealed that this station matched mesopelagic environments more closely than surface waters or deep chlorophyll maxima. Research from Farrenkopf *et al*. indicated that bacteria are responsible for IO_3_^-^ reduction in oxygen minimum zones^12,65^. Further, Saunders *et al*. showed a preferential expression of AioA-like genes in the Eastern Pacific oxygen minimum zones, which our evidence now suggests are IO_3_^-^-reductases (IdrA)^30^.

To test whether locations with *idrA* possessed a unique chemical signature, we ran a principal component analysis using the variables associated with sample environments. Together the first two components of these geochemical variables explained 70.66% of the variance observed between *idrA* present and *idrA* absent samples. We determined that *idrA* presence was correlated most strongly with increased nitrate, phosphate, and silicate concentrations (Figure 6C-E). Additionally, *idrA* presence was negatively correlated with dissolved oxygen concentrations (Figure 6C-E). Such an observation is atypical for highly productive nitrate and phosphate depleted OMZs^60,66,67^. A possible explanation for this observation is that DIRM inhabit a unique niche above OMZs where residual O_2_ prevents *fnr*-dependent expression of nitrate reductase^68^. Organisms in these environments could potentially use IO_3_^-^ as an alternative electron acceptor. Excess phosphorous in these zones seemingly serves as a proxy indicator of lower overall productivity, and potentially reflects the limiting concentration of IO_3_^-^ and oxygen for biomass accumilation^4,23^. Our explanation corroborates results from Farrenkopf *et al*. that shows an I^-^ maximum occurring at the boundary of the OMZ^61^, but further studies into the biochemistry of IO_3_^-^ reduction under suboxic conditions and the contribution of DIRM to I^-^ formation at this transition zone are necessary to undeniably link the I^-^ maximum with the presence of *idrA* directly.

### Significance

Here we describe a new organism, *Denitromonas iodocrescerans*, that grows by IO_3_^-^ respiration which is mediated by a novel molybdenum dependent DMSO reductase. The conserved core genes associated with DIR and the chemistry of iodine oxyanions are consistent with a hybrid enzymatic-abiotic pathway by which IdrAB reduces IO_3_^-^ to HIO, which abiotically disproportionates to I^-^ and IO_3_^-^^16,37^. In this model, cytochrome c peroxidase like proteins (IdrP_1_ and IdrP_2_) detoxify reactive H_2_O_2_ byproducts. Genes for this metabolism are part of a highly conserved IO_3_^-^ reduction genomic island (IRI). Organisms harboring the IRI belong to phylogenetically distinct taxa, many of which are associated with marine sediments or multicellular hosts, suggesting that DIR is a horizontally transferred metabolism across marine ecosystems over geologic time. The abundance of IdrA genes across ocean metagenomes strongly correlates to oxygen minimum zones, indicating a niche for this metabolism in low-oxygen, high nitrate habitats across the ocean, from sediments to oxygen-minimum zones to the surfaces of multicellular organisms. In high-nitrate, low-oxygen conditions, bacteria with the IRI can use IO_3_^-^ as an electron acceptor to obtain energy from the oxidation of organic matter. IO_3_^-^ is constantly replenished by the chemical oxidation of I^-^, so DIRM do not rely on other organisms for their substrate. IO_3_^-^ is typically scarce (0.45μM in seawater)^4^, so DIRM must compete with IO_3_^-^ reduction by chemical reductants and by inadvertent biological activity, such as by algae, that contribute to the relative depletion of IO_3_^-^ in those waters^7,61,65,69,70^. By analogy, perchlorate-reducing bacteria, which are common but sparse due to low natural abundance of perchlorate^71^, may provide further insight into the ecology of DIRM broadly. The rarity of IO_3_^-^ reduction genes among bacteria despite the ability of the metabolism to be horizontally transferred likely reflects the evolutionary constrains of growth by DIR. Intriguingly, one organism, *Sedimenticola thiotaurini*, seemingly possesses both perchlorate and IO_3_^-^ reduction pathways, presenting future opportunities to study the ecology of these metabolically versatile microorganisms^72^. Moreover, organisms such as *Vibrio spp*. and *Moritella spp*. show some degree of vertical transfer for the IRI throughout recent evolutionary history, indicating possible niches among sea fauna and cold environments where DIR is biogeochemically favorable. Future studies addressing the affinity of IdrAB for IO_3_^-^ may also shed light on how DIRM thrive at such low environmental concentrations. Additionally, further research into the chemistry of iodine oxyanions may provide insight on the intermediates of IO_3_^-^ reduction. Addressing these open questions may ultimately shed light on new potential niches for DIRM and provide a role for these organisms in potentiating iodine redox cycling globally.

### Description and Phylogeny of *Denitromonas iodocrescerans* sp. nov. strain IR-12^T^

*Denitromonas iodocrescerans* (i.o.do.cre’scer.ans) Chem. n. *iodo* as it pertains to iodine; L. pres. part. *crescerans* for growing; N.L. pres. part. *iodocrescerans* iodine-growing.

*D. iodocrescerans* is a facultatively anaerobic chemoorganotroph, gram negative, rod-shaped, 1.5-2.0 μM long by 0.6-0.7 μM wide, and motile by means of a unipolar flagellum (Figure 1B). Colonies are circular, smooth, and range in color from transparent to an opaque/whitish-sky blue color after 48 hours of growth on R2A agar at 30°C. Extended growth on R2A agar (96 or more hours) results in a light coral pink colony color. *D. iodocrescerans* grows by oxidizing D-glucose, lactate, or acetate with concomitant reduction of oxygen (O2), nitrate (NO3^-^), or iodate (IO_3_^-^). It grows on up to 4 mM of iodate with an optimum at 2 mM. Additionally, the organism can tolerate up to 6.25 mM of iodide. Growth occurs between 20-30°C with an optimum of 30°C. It grows at a range of 0-5% salinity with an optimum of 3% NaCl on minimal media. *D. iodocrescerans* has an innate resistance to tetracycline (10 μg/μL) and chloramphenicol (25 μg/μL) but is sensitive to kanamycin, which inhibits growth at concentrations as low as 5 μg/μL.

The genome of *D. iodocrescerans* is 5,181,847 bp (average coverage 64.2x) with 4697 CDS, a G+C content of 66.54%, 57 tRNAs, one tmRNA, one CRISPR, and a single plasmid 81,584 bp long whose function remains unclear. The full genome has been deposited in GenBank (BioProject ID PRJNA683738) currently consisting of 202 contigs. Phylogenetically, *D. iodocrescerans* belongs to the class *Betaproteobacteria*; however, its phylogeny beyond this class becomes less clear. The 16S rRNA locus suggests that *D. iodocrescerans* is a subclade of *Azoarcus*, which belongs to the family *Zoogloeaceae*^73^. However, the NCBI database suggests that the genus *Denitromonas* belongs to the family *Rhodocyclaceae*.

The type strain of *Denitromonas iodocrescerans*, IR-12^T^ was enriched from marine sediment from the Berkeley Marina in the San Francisco Bay during the Fall of 2018 (further details explained in methods below). The strain has been deposited in the American Type Culture Collection (ATCC XXXXX).

## Acknowledgements

The authors acknowledge Mariana Shalit, Dylan Dang, Jessica Kretschmer, Rachael Peng, Mitchell Thompson, and Hans Carlson for lab support and advice throughout the project. Funding for research on iodate in the Coates lab was provided to VRU through the NSF GRFP Base Award: DGE1752814.

## Contributions

JDC guided the research. VRU and KL performed all physiology experiments and measurements. VRU performed all cloning experiments. VRU and TPB performed the comparative genomic analysis and phylogenetic analyses. VRU and ZH performed the analysis of the TARA Oceans data. VRU and JDC developed the model. VRU wrote the draft manuscript and created the figures with guidance from JDC. All authors contributed to data analysis, reviewed the manuscript, and approved of its publication.

## Methods

### Media, chemicals, and culture conditions

Anaerobic enrichment cultures from marine environments were grown at 30°C using a minimal media containing the following per liter: 0.54g NH_4_Cl, 0.14g KH_2_PO_4_, 0.20g MgCl_2_ · 6 H_2_O, 0.14g Na_2_SO_4_ · 10 H_2_O, 20.0g NaCl, 0.24g Na_2_MoO_4_ 0.20g, and 2.5g NaHCO_3_ with an added vitamin mix and mineral mix. Oxygen was removed from the media and bottles were dispensed in an 80%N_2_/20%CO_2_ atmosphere. Anaerobic subcultures for isolation were grown in Artificial Pore Water (APM) medium at 30°C (30.8g NaCl, 1.0g NH_4_Cl, 0.77g KCl, 0.1g KH_2_PO_4_, 0.20g MgSO_4_·7H_2_O, 0.02g CaCl_2_ · 2 H_2_O, 7.16g HEPES, along with vitamin and mineral mixes. A post sterile addition of 34.24mL 0.4M CaCl_2_ and 26.07mL 2M MgCl_2_ · 6H_2_O was added to all APM media. Conditions with lactate, acetate, iodate, and nitrate all used the sodium salts of these compounds. Conditions without molybdenum omitted Na_2_MoO_4_ from the mineral mixes. Aerobic cultures were all grown either on APM, R2A (HiMedia, USA), or R2A agar (BD Biosciences, USA). Kanamycin concentrations when used were at one tenth the standard concentrations on plates (5 mg/L, Sigma Aldrich, USA) and at one fourth the standard concentration in liquid (12.5 mg/L). All compounds were purchased through Sigma Aldrich (Sigma Aldrich, USA). Growth of tubes were measured either using the Thermo Scientific^™^ GENESYS^™^ 20 or the TECAN Sunrise^™^ 96-well microplate reader set at a wavelength of 600 nm. For growth measurements in Hungate tubes, a special adapter was built to measure the tubes on the GENESYS™ 20. Growth experiments using the microplate reader were run in an anerobic glove bag.

### Isolation of dissimilatory iodate-reducing bacteria

Sediment from the oxic/anoxic boundary layer in the San Francisco Bay estuary (37°86’56.4” N, −122°30’63.9” W) was added to anaerobic media bottles at 25g/100mL for isolation of dissimilatory iodate-reducing bacteria. Samples were degassed and amended with acetate and iodate to enable growth of heterotrophic iodate reducing bacteria. Enrichments that showed iodate reduction to iodide were then passaged at least five times into fresh minimal media with 10mM acetate and 2mM iodate. To ensure purity of the passaged enrichment culture, the organism was plated aerobically onto an agar plate containing the minimal media, and a single colony was isolated from this plate.

### Strains and plasmids

All plasmids, primers and strains constructed are listed in Table S1. The *E. coli* strain used for plasmid propagation was XL1-Blue, while WM3064 was used to perform conjugations. Plasmid pNTPS138, a generous gift from the Kathleen Ryan Lab at UC Berkeley, was used for the SacB counterselection. Plasmid pBBR1-MCS2 is a low copy expression vector and was used for complementation experiments. All expression plasmids and deletion vectors were constructed using the Benchling software suite (San Francisco, USA). Plasmids were assembled either by Gibson assembly or restriction digestion and ligation using standard procedures. Gibson assembly was carried out using NEB HiFi 2x Master Mix, and remaining enzymes and master mixes were ordered from New England Biosciences (NEB, USA). Plasmids were routinely isolated using the Qiaprep Spin Miniprep kit (Qiagen, USA), and all primers were ordered from Integrated DNA Technologies (IDT, Coralville, IA). Amplification of DNA for generating assembly products was performed using Q5 DNA Polymerase 2x Master Mix (NEB, USA) with 3% DMSO. Amplification of distinct portions of the genome were optimized since most sequences in the iodate reduction cluster contain at minimum 60% GC content, making amplification relatively challenging. All *D. iodocrescerans* strains (pre- or post-transformation) were propagated from glycerol stocks (25% glycerol) stored at −80°C, grown on a plate for up to 72 hours, picked and then grown for an additional 48-72 hours in liquid R2A. For additional information on performing transformations and conjugations in *D. iodocrescerans* see supplemental methods.

### Iodate and iodide quantification

A Dionex^™^ IonPac^™^ AS25 Anion Exchange Column (Thermo Fischer, USA) was used exclusively to measure the consumption of iodate and acetate, as well as the production of iodide in all samples. Briefly, all samples are diluted 1:25 in deionized water and loaded onto the autosampler for processing. Standards are made by serial dilution starting with 1 mM of the standard molecule. All samples were run in triplicate. Acetate peaks were consistently detected at 3.6 minutes, iodate peaks were consistently detected at 3.8 minutes, and iodide peaks were consistently detected at 11.5 minutes at a flow rate of 1mL/min.

### Genome sequencing, comparative genomics, and phylogenetic analysis

Genome sequencing was carried out on an Illumina HiSeq4000 using 150bp paired end reads. The genome was subsequently assembled using SPAdes 3.9^74^ and the assembly graph was assessed for completion using bandage^75^. The Prokka (version 1.14) pipeline was then used to generate the genome annotations and the general feature format file (.gff), which allowed for genome navigation and visualization on the Artemis software (available at http://sanger-pathogens.github.io)^76^. To search for the iodate reduction island, MMseqs2 was used to cluster homologous proteins in the amino acid FASTA (.faa) files from *D. iodocrescerans, P. stutzeri* sp. SCT, *D. halophilus* SFB-1, and *P. stutzeri* sp. CAL by subfamily^41^. A presence and absence matrix for each subfamily was generated and represented as a four-way Venn diagram using pyvenn (https://github.com/tctianchi/pyvenn). To identify additional iodate reductase proteins in public databases, a profile-HMM was constructed using HMMER 3.0 following a multiple sequence alignment using MUSCLE 3.8 on the molybdopterin oxidoreductase (Pfam_00384) seed set and *D*. iodocrescerans/*P. stutzeri* SCT IdrA proteins^77,78^. A separate arsenite oxidase (AioA) profile-HMM was created using analogous methods. Genomes from high probability HMM hits (threshold above 640 on https://www.ebi.ac.uk/Tools/hmmer/search/phmmer) and BLAST hits were downloaded from NCBI using ncbi-genome-download (https://github.com/kblin/ncbi-genome-download). Approximately-maximum-likelihood phylogenetic trees were generated using Fasttree^79^ specifying 10,000 resamples and using standard settings for everything else. Visualization of resultant trees used the ete3 toolkit^80^. To perform the neighborhood frequency analysis, 10 genes upstream and downstream from the *aioA* or *idrA* locus were extracted from the associated GenBank files for each genome, and MMseqs2 was used to cluster homologous proteins into subfamilies ^41^. To search for cld in the downloaded genomes, a profile-HMM for cld, described previously, was used^81^. Frequency was calculated as number of genomes in possession of a cluster divided by the total number of genomes. Projections of this data were drawn using a custom Python 3.7 script. All tanglegram analyses used Dendroscope to load trees for processing and visualization^47^.

### Distribution of iodate reductase in ocean metagenomes

The profile-HMM for iodate reductase (described above) was used to search all 40 million non-redundant open reading frames from the 243-sample Tara oceans dataset. Open reading frames were downloaded (available from https://www.ebi.ac.uk/ena/data/view/PRJEB7988) and translated to amino acid sequences using custom BioPython code^82,83,84^. The amino acid sequences in the 0.22-micron and 0.45-micron range were then searched for hits using the IdrA profile-HMM set at a threshold score of 640. Hits were then grouped by station for further analysis. Reads were mapped to scaffolds with Bowtie2^85^ and reads were counted using SAMtools^86^. Read abundance mapping to these unique IdrA hits were quantified by using the transcripts per million (TPM) method for read quantification as described in Ribicic *et al*^58,59^. Ten variables in the metadata associated with the chemical environment at each sampling location were analyzed using the principal component analysis module on scikit-learn 0.23.1^87^ All sites regardless of *idrA* presence were included in the analysis. Missing metadata values were imputed using the Multivariate Imputation by Chained Equations method (MICE)^88^. Variables included in the analysis were ‘Sampling depth [m]’, ‘Mean_Temperature [deg C]’, ‘Mean_Salinity [PSU]’,’Mean_Oxygen [umol/kg]’, ‘Mean_Nitrates[umol/L]’, ‘NO2 [umol/L]’, ‘PO4 [umol/L]’, ‘SI [umol/L]’,’NO2NO3 [umol/L]’, and irradiance ‘AMODIS:PAR8d,Einsteins/m-2/d-1’. Components were built using “pca.fit_transform()” and confidence ellipses at one standard deviation were set for each group. Component coefficients were extracted from principal components by using “pca.components_” and displayed as a loadings plot. Explained variance was also extracted from “pca.components_” to display on PCA axes. The map of *idrA* abundance was created using Cartopy 0.17.

