## Supplementary figures and images for "Genetic and phylogenetic analysis of dissimilatory iodate-reducing bacteria identifies potential niches across the world’s oceans"

### Fig S1

Aldr Avs Rps3

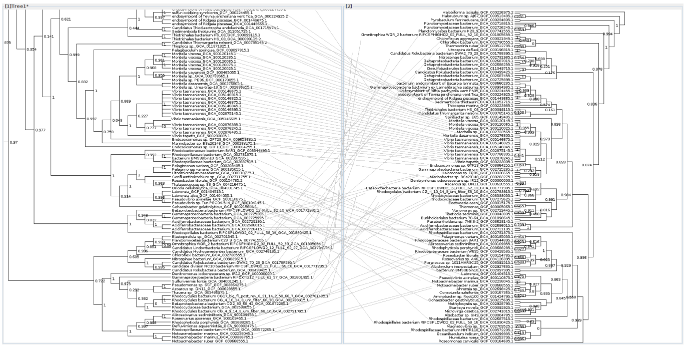

C IdrP1/IdrP2 vs Rps3

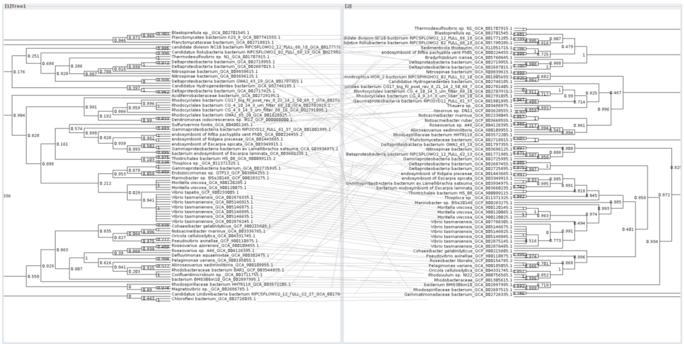

B IdrB vs Rps3

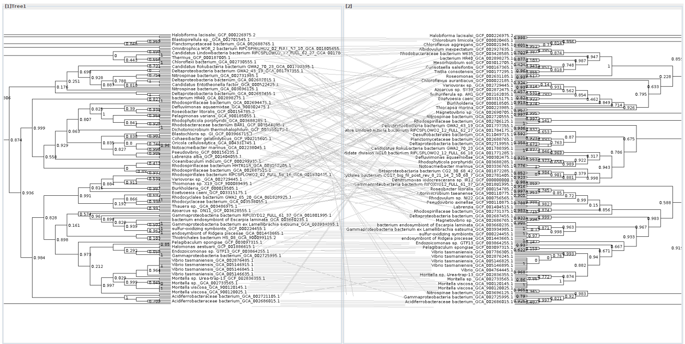

D IdrAvs IdrB

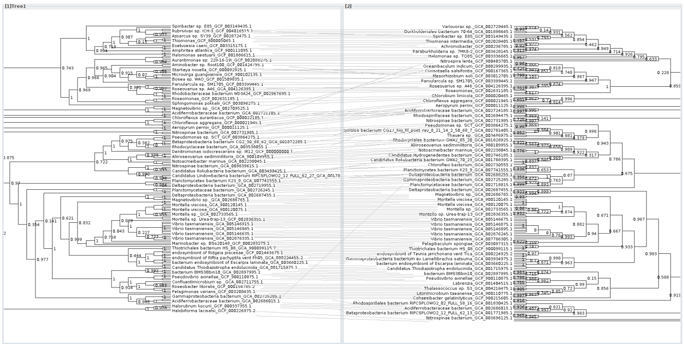

### Fig S2

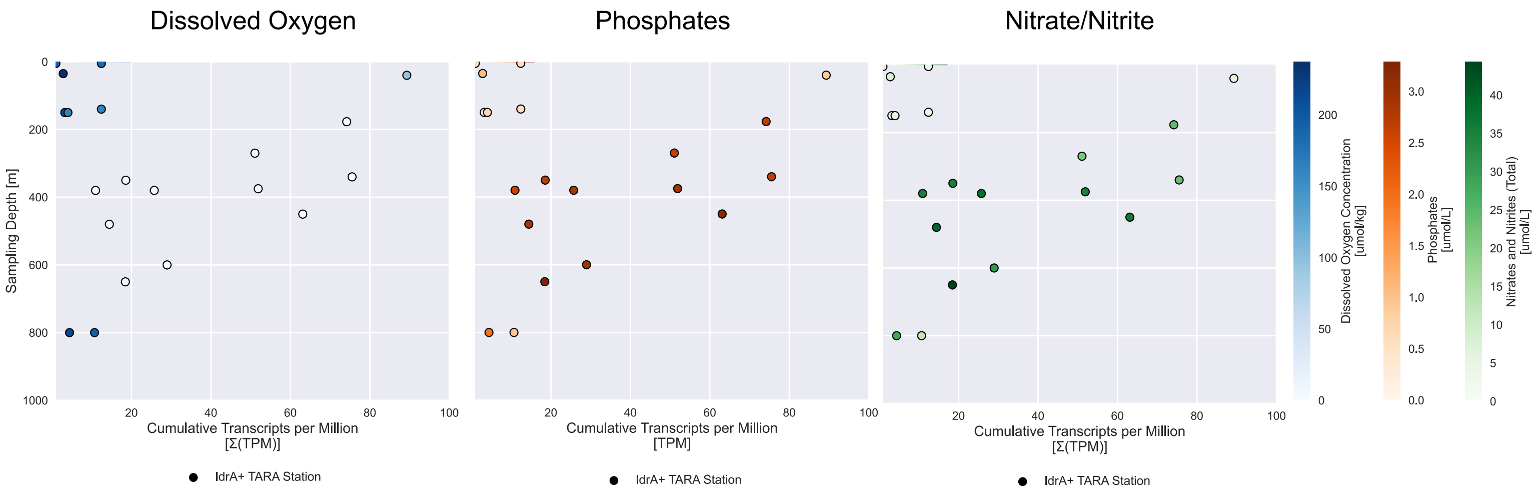
